# S-PLM: Structure-aware Protein Language Model via Contrastive Learning between Sequence and Structure

**DOI:** 10.1101/2023.08.06.552203

**Authors:** Duolin Wang, Mahdi Pourmirzaei, Usman L Abbas, Shuai Zeng, Negin Manshour, Farzaneh Esmaili, Biplab Poudel, Yuexu Jiang, Qing Shao, Jin Chen, Dong Xu

## Abstract

Proteins play an essential role in various biological and engineering processes. Large protein language models (PLMs) present excellent potential to reshape protein research by accelerating the determination of protein function and the design of proteins with the desired functions. The prediction and design capacity of PLMs relies on the representation gained from the protein sequences. However, the lack of crucial 3D structure information in most PLMs restricts the prediction capacity of PLMs in various applications, especially those heavily dependent on 3D structures. To address this issue, we introduce S-PLM, a 3D structure-aware PLM that utilizes multi-view contrastive learning to align the sequence and 3D structure of a protein in a coordinated latent space. S-PLM applies Swin-Transformer on AlphaFold-predicted protein structures to embed the structural information and fuses it into sequence-based embedding from ESM2. Additionally, we provide a library of lightweight tuning tools to adapt S-PLM for diverse protein property prediction tasks. Our results demonstrate S-PLM’s superior performance over sequence-only PLMs on all protein clustering and classification tasks, achieving competitiveness comparable to state-of-the-art methods requiring both sequence and structure inputs. S-PLM and its lightweight tuning tools are available at https://github.com/duolinwang/S-PLM/.

## Introduction

Proteins play a crucial role in resolving many of the health, energy, and environmental challenges facing society today. An essential task is to gain information about proteins of interest quickly and accurately. Along this line, computational predictions of protein properties from protein sequences play important roles. Protein language models (PLMs) can reveal the underlying patterns within protein sequences and predict properties based on the underlying sequence patterns^1^. The current paradigm for PLM development and deployment includes two stages: (1) train a PLM to convert the amino acid sequence into a mathematical representation (embedding) by means of the masked language modeling (MLM) or autoregressive strategy, where the model predicts masked or next amino acids in the sequence based on the surrounding or previous context^2–4^; and (2) adapt the pretrained PLM with the protein property^2^ data to perform specific protein tasks. Several PLMs have been developed following this paradigm, such as the ProtBert^3^, ESM2^5^ models and ProtGPT2^6^. These PLMs have shown encouraging results for protein property predictions and *de novo* protein designs and demonstrated their potential for new knowledge discovery and analyses^7–9^.

One challenge in developing PLMs is enriching critical biophysical information into the embeddings. It is well known that a protein’s function relies on its 3D structure. However, most PLMs are trained solely on amino acid sequences, thereby constraining their predictive capabilities, especially those heavily depending on 3D protein structures. Some methods have been developed to enrich the evolutionary or functional information in the sequence-based embedding. One method integrates multiple sequence alignments (MSAs) into PLMs, such as AlphaFold’s Evoformer^10^ and MSA Transformer^11^. ProteinBERT incorporated gene ontology (GO) annotations into the MLM pretraining scheme^12^. It utilized a denoising autoencoder to pretrain with corrupted protein sequences and GO annotations and performed comparatively well in many protein tasks despite its relatively smaller size. These methods enriched the information in the sequence-based embedding and boosted the performance of the PLMs. However, none of these methods incorporated the critical structural information in an indirect manner.

Another emerging method is to develop a joint embedding with both sequence and structural inputs. Chen et al. proposed a self-supervised learning-based method for protein structure representation to leverage the available pretrained PLMs^13^. Zhang et al. systematically explored the joint embedding for proteins based on ESM2 and three distinct structure encoders^14^. Hu et al. also developed a joint embedding for proteins by coupling protein sequence and structure^15^. These joint embeddings performed better in numerous protein property prediction tasks, highlighting the importance of including structural information in the protein representation. However, the utilization of these joint-embedding models requires both sequence and structure as input. Even with prediction tools such as AlphaFold, gaining reliable structures for some specific proteins remains a challenge. Furthermore, computationally, these methods take an extra step to obtain predicted structures, which requires extra time and effort.

We propose an alternative approach to protein representation learning. Instead of creating a joint embedding that needs both sequence and structural inputs, we have developed a sequence-based embedding that incorporates structural information. One feasible strategy for incorporating structural information into sequence-based embeddings is through cross-view representation learning. PromptProtein^16^ exemplifies this approach by pretraining on a sequence-to-structure prediction task to generate structure-aware representations. In contrast, our approach is based on another technique known as multi-view contrastive learning. Unlike cross-view representation learning, which translates embeddings from one view to another, multi-view contrastive learning aligns embeddings into a coordinated latent space across multiple views. This technique has garnered significant attention for its ability to capture rich and complementary features from diverse perspectives^17,18^. During training, the contrastive loss function pushes the representations of similar views (sequence and structure of the same protein) to be close to each other in the embedding space while simultaneously separating representations of dissimilar views (between different proteins) further apart. As a result, multi-view contrastive learning effectively captures underlying semantic patterns in the data. A head-to-head comparison has demonstrated that multi-view contrastive learning can be more effective than cross-view representation learning^18^.

To this end, we propose S-PLM, a 3D structure-aware PLM that enables the sequence-based embedding to carry the structural information through multi-view contrastive learning. One advantage of S-PLM compared to those joint-embedding models is that after training, S-PLM only needs amino acid sequences as the input during inference. This sequence-only input avoids the overhead of using predicted protein structures. It is worth mentioning that two other recent successful cases have illustrated the potential of contrastive learning in enriching information in sequence-based embedding. One is the ProtST, which injected biomedical text into a PLM by aligning these two modalities through a contrastive loss^19^. After training, their PLM contained enhanced protein property information and demonstrated superiority over previous models on diverse protein representations and classification benchmarks. Another one is the CLEAN^20^ model, which utilized information from the EC number to enhance the ability of PLM to predict enzymatic functions based on protein sequence. Although S-PLM can accommodate either structure or sequence as the input, this paper will only demonstrate the advantages of S-PLM in encoding protein sequences and leveraging structural information to improve sequence representations for various protein prediction tasks. The sequence encoder of S-PLM was implemented based on a pretrained ESM2 model, offering extensibility to incorporate new protein properties incrementally without forgetting previous knowledge in the model^21^. Through several unsupervised protein clustering tasks and several structure-dominated supervised protein prediction tasks, S-PLM outperformed the baseline ESM2 model significantly.

To apply pretrained PLMs to specific protein prediction tasks, a typical domain adaptation approach involves fine-tuning the PLMs on domain-specific data to update all model parameters. However, full fine-tuning may not work well for large PLMs due to limited domain-specific training data that may lead to catastrophic forgetting^22^ or severe computational and memory costs. To address this challenge, researchers have developed a set of lightweight tuning strategies, making specific and parameter-efficient modifications to large preexisting models. These strategies, such as fine-tune top layers, adapter tuning^21^, and low-rank adaptation (LoRA)^23^, selectively update targeted parameters while keeping others frozen, significantly reducing computational and memory requirements and mitigating data scarcity while achieving comparable or superior performance. However, there is limited existing research on applying these methods to PLMs^24,25^. In this work, we explored several lightweight tuning strategies for diverse protein prediction tasks utilizing S-PLM. Through further lightweight tuning, S-PLM achieved the best performance in certain categories of gene ontology^26^ (biological process and molecular function) and protein secondary structure prediction tasks. While it exhibited suboptimal yet comparable performance to cutting-edge methods employing both sequence and structure inputs in cellular components of gene ontology and enzyme commission number^27^ prediction tasks. A library of lightweight tuning methods has been made available at https://github.com/duolinwang/S-PLM/.

## Results

### Structure-aware protein language model (S-PLM)

Figure 1a depicts the framework of S-PLM. The pretraining architecture of S-PLM consists of two encoders, one to encode protein sequences and the other to encode 3D protein structures. In this study, the one-letter amino acid sequences are utilized as the input of protein sequences. The backbone C_α_ contact maps are used to represent the protein 3D structures because inter-residue distances contain comprehensive and essential information about protein structure. During pretraining, S-PLM inputs both amino acid sequences and backbone C_α_ contact maps. The protein sequence information is converted into the amino-acid level (AA-level) embedding through a sequence encoder (ESM2-Adapter), while the contact map information is transformed into a protein-level embedding through a structure encoder (Swin-Transformer^28^). Then through each corresponding projector layer, sequences and contact maps are converted into separate protein-level embeddings (Methods 1 and 2). Finally, the S-PLM model is trained using contrastive learning to minimize the contrastive loss for a batch of sequences and contact maps. The objective of the S-PLM model is to maximize the alignment of the embeddings for the sequence and structure from the same protein and clearly separate the embeddings for the sequences and structures between different proteins (de-alignment). Inspired by the SimCLR method^29^, our work adapts the CLIP^30^ approach for contrastive language-image pretraining. Besides the CLIP’s alignment and de-alignment across different modalities, our model also accounts for de-alignment within the same modality. For instance, as shown in Figure 1a, our model also emphasizes dissimilarity between embeddings of sequences (*s*_1_ ↔ *s*_2_) and embeddings of contact maps (*c*_1_ ↔ *c*_2_) from different proteins (Methods 3).

**Figure 1.**
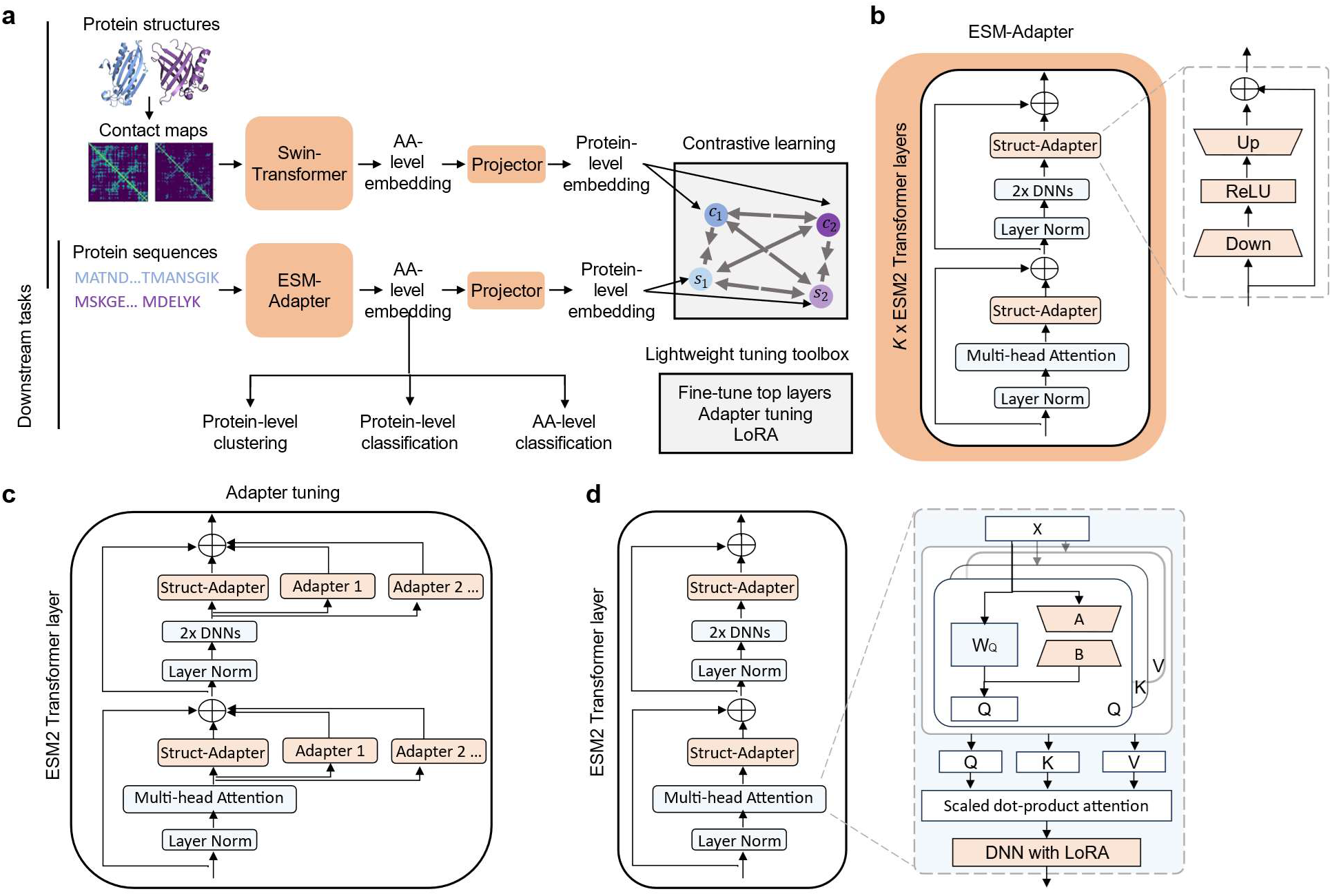
The framework of S-PLM and lightweight tuning strategies for downstream supervised learning. ***a***, The framework of S-PLM. During pretraining, the model inputs the amino acid sequences and contact maps simultaneously. The protein sequences are transformed into AA-level embeddings through an ESM-Adapter, while the contact maps are transformed into protein-level embeddings using Swin-Transformer. With respective projector layers, the S-PLM model is trained through contrastive learning on the protein-level embeddings from each modality. After pretraining, the ESM-Adapter that generates the AA-level embeddings before the projector layer is used for downstream tasks, such as protein-level clustering, classification, and AA-level classification tasks. The entire ESM-Adapter model can be fully frozen or learnable through lightweight tuning. ***b***, Architecture of the ESM-Adapter. The sequence encoder of S-PLM (ESM-Adapter) is tuned on the top-*K* Transformer layers of ESM2, each of which adds a compact adapter module (Struct-Adapter). During the pretraining, the ESM2 backbone model is frozen, and only the Struct-Adapter modules are trainable. ***c***, Adapter tunning for supervised downstream tasks is implemented by integrating a list of additional paralleled adapters on the Struct-Adapter modules. ***d***, LoRA tuning for supervised downstream tasks is implemented by adding trainable rank decomposition matrices into the multi-head attention layer of top-*K* Transformer layers of ESM2.

The sequence encoder of S-PLM was implemented as a structure adapter on the ESM2 model using adapter tuning^21^. Our adapter tuning involves integrating adapter modules into the top-*K* Transformer layer of the ESM2 model. As shown in Figure 1b, the adapter modules (Struct-Adapter) consist of a bottleneck structure and a skip-connection, positioned twice in one Transformer layer: after the multi-head attention projection and after the two feed-forward layers (Methods 5). Using adapter tuning to implement the sequence encoder of S-PLM offers several advantages. First, the integrated adapter module is compact. It contains far fewer parameters than the original Transformer modules of ESM2, which alleviates the training burden. Second, it allows for continuous training to add new protein features for the future extension of our model, such as to include protein function, without catastrophic forgetting of previously learned features, because the S-PLM pretraining retains the sequence representation capabilities of the ESM2 model with the entire ESM2 backbone model intact.

During the inference stage, S-PLM has the flexibility to accept either sequence or contact map as the input and produces corresponding embeddings at various levels tailored to specific downstream tasks (e.g., protein-level clustering, protein-level classification, and AA-level classification). This versatility allows S-PLM to adapt and provide suitable representations based on the specific input data and requirements of the given task. In this paper and the subsequent results, the S-PLM model primarily generates sequence embeddings from protein sequences. Therefore, the pre-trained sequence encoder of S-PLM that generates the AA-level embeddings before deploying projectors is used for downstream tasks. The entire sequence encoder can be fully frozen or learnable. To fully exploit the potential of S-PLM in supervised protein prediction tasks, we have developed several lightweight tuning strategies based on the sequence encoder of S-PLM, all of which are incorporated into the lightweight tuning toolbox, including the fine-tune top layers, adapter tuning (Figure 1c), and LoRA tuning (Figure 1d) (Methods 5).

### Contrastive learning rearranges the alignment between the sequence and structure embeddings

We investigated the impact of contrastive learning on the alignment between sequence and structure embeddings. Initially, we randomly selected 25 proteins from an independent test set (500 proteins in total) and projected their corresponding sequence and structure embeddings onto a 2D t-SNE space. We employed trajectory plots of sequence and structure embeddings to illustrate the movement of embeddings during contrastive learning. As depicted in Figure 2a, before contrastive learning, the sequence-based embeddings (obtained from pretrained ESM2-t33_650M_UR50D) and the structure-based embeddings (obtained from pretrained Swin-Transformer) were distinctly distributed in the 2D t-SNE space. Embeddings from the same modality were closer than those from different modalities, indicating no information exchange between the two modalities. Conversely, after contrastive learning, sequence and structure embeddings of the same protein moved closer, whereas those of different proteins became more separated, suggesting successful integration of protein structure information into sequence-based embeddings (and vice versa). To show the effect of protein size in our training strategy, we grouped the proteins in the test dataset into different bins based on their sequence length. Figure S1 (Supporting Information) shows the embedding distributions after contrastive learning within specific sequence length ranges. This experiment shows that the observed alignment is not influenced by sequence length.

**Figure 2.**
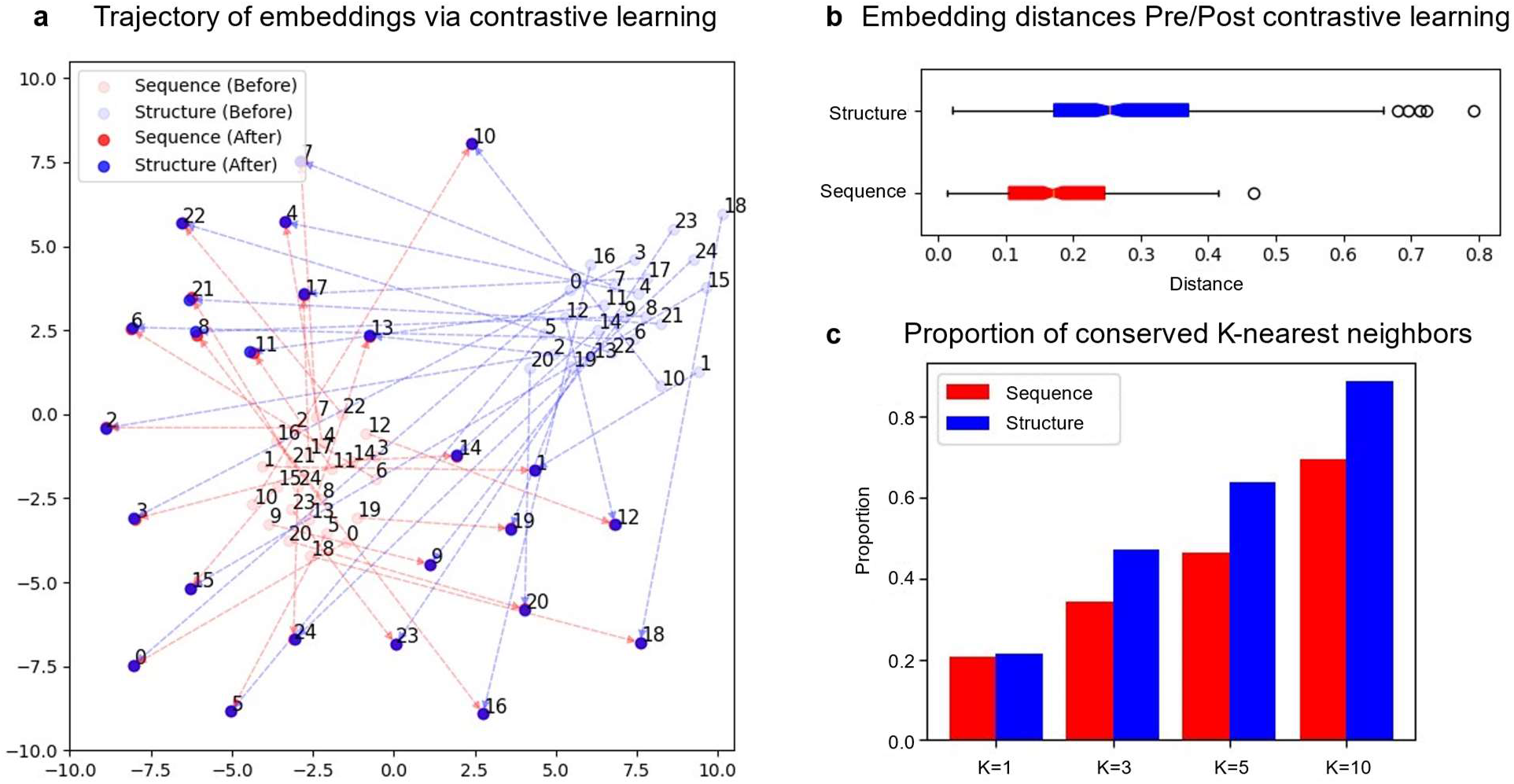
Structure and sequence embedding rearrangement through contrastive learning. **a**, Trajectories illustrating the alignment of sequence and structure embeddings via contrastive learning. A blue node indicates a structure-based 2D-t-SNE embedding for a protein, whereas a red node indicates a sequence-based 2D-t-SNE embedding for a protein. The sequence and structure embeddings of the same protein after contrastive learning are almost identical such that their solid circles overlap. The number beside each node indicates the protein index (from 0 to 24). **b**, Distances between sequence (structure) embeddings before and after contrastive learning. The distances are calculated by the Euclidean distance on the original sequence (structure) embedding space. **c**, Proportion of conserved *K*-nearest neighbors. Each bar represents the proportion of conserved *K*-nearest neighbors for a specific value of *K*. Five hundred proteins were used for the analyses in panels b and c.

Notably, the changes in structure embeddings were more pronounced than those in sequence embeddings, causing them to spread around the original embeddings. This observation is further supported by Figure 2b, which shows the distribution of embedding distances for each modality across the entire 500 testing proteins before and after contrastive learning, revealing more significant embedding changes in structure than in sequence. This phenomenon also underscores that our sequence encoder of S-PLM effectively retained the sequence representation capabilities of the base ESM2 model, minimizing catastrophic forgetting. Additionally, we quantitatively assessed the effectiveness of contrastive learning in preserving the neighborhood topology and relationships within sequence and structure embeddings by calculating the proportion of conserved K-nearest neighbors for each modality after contrastive learning across different values of *K* (Figure 2c). For the entire set of 500 testing proteins, over 20% of the first-nearest neighbors of both the sequence and structure embeddings were preserved after contrastive learning. Increasing *K* to 10 preserved 69% of sequence and 89% of structure embedding neighbors. Across all *K* values, the proportions of preserved structure embedding neighbors were higher than those of sequence, indicating that our training process effectively preserved more topological and geometrical information within the structure embeddings.

### S-PLM injects the structural information into the sequence latent space

To investigate whether S-PLM can inject the structural information into the sequence latent space, we evaluated the sequence representations for the CATH protein domains^31^. Because this experiment prioritizes structural information, we deliberately chose a single representative sequence from each CATH superfamily to provide clear visualization. We utilized the CATHS40 dataset, whose proteins have maximal 40% sequence identity. Our analysis focused exclusively on the class, architecture, and topology levels of the CATH hierarchy, excluding the homologous superfamily at the last level, which comprises clusters primarily driven by sequence similarity. We visualized and benchmarked the sequence representations produced by S-PLM against models that rely solely on sequence information, including ESM2 (ESM2_t33_650M_UR50D); the pretrained PLM on which S-PLM is based; and PromptProtein^16^, another structure-aware model that utilizes structure prompts pretrained by predicting 3D structures from sequences (Figure 3a). Each row of Figure 3a shows the t-SNE visualization of protein embeddings from the five most represented categories of one hierarchy. It shows that sequence representation from S-PLM separates CATH structural classes more clearly than the embeddings from the other two models.

**Figure 3.**
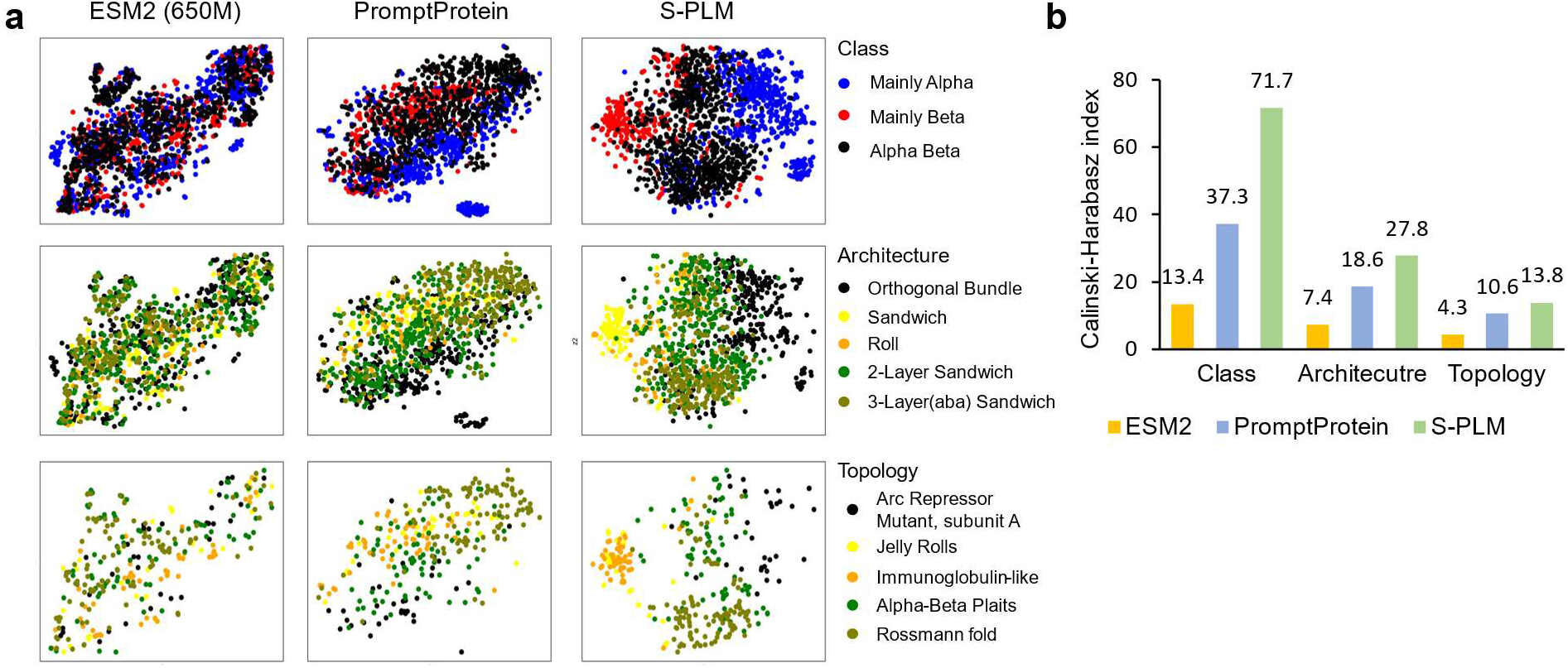
Visualization and benchmark of protein embeddings for three CATH structural hierarchies (class, architecture, and topology). **a**, t-SNE visualization of protein embeddings from the five most represented categories from one hierarchy. Embeddings produced by ESM2 and PromptProtein are shown side by side with S-PLM. **b**, Utilizing the CHI to quantitatively assess the capability of embeddings derived from different methods in clustering CATH structural categories.

We further utilized the Calinski-Harabasz index (CHI)^32^ to quantitatively assess the capability of embeddings derived from different methods in distinguishing CATH structural categories. The CHI score quantifies the ratio between the sum of between-cluster dispersion and the sum of within-cluster dispersion. We applied the CATH categories to define ground-truth clusters, using sequence embeddings to calculate both between- and within-cluster dispersion. As shown in Figure 3b, S-PLM achieved much higher CHIs across all CATH hierarchy levels than ESM2 and PromptProtein. Given that these CATH categories were established using protein structures, this analysis strongly suggests that the sequence embedding produced by the developed S-PLM exhibits an inherent awareness of protein structures, surpassing the other two PLMs in effectively distinguishing proteins with diverse structural characteristics. The poor performance of the sequence-only ESM2 also indicated its limitations in explicitly acquiring protein structure knowledge.

### Clustering enzymes via S-PLM

Recently, Huang et al. introduced a structure-based protein clustering approach for discovering deaminase functions and identifying novel deaminase families^33^. They applied AlphaFold2 to predict protein structures and subsequently clustered the entire deaminase protein family based on the predicted structure similarities through structure alignment. They discovered new functions of the deaminase proteins and new deaminases; such findings cannot be obtained by mining amino acid sequences. In this study, we investigated the effectiveness of our S-PLM model in clustering the deaminase family by comparing and benchmarking the sequence embeddings from S-PLM against two other PLMs (i.e., the ESM2 and PromptProtein). We utilized the same sequence data as Huang et al. to generate representations for each query protein sequence. For this task, we utilized the S-PLM and obtained a 1280-dimensional embedding for each protein. Subsequently, we employed t-SNE to reduce the vector to a 2D representation and applied the K-Means clustering method to the reduced dimension. The Adjusted Rand Index (ARI) was computed by comparing K-Means clustering assignments with known deaminase family annotations. We conducted the t-SNE visualization and calculated the ARI in the same way for all the benchmarked methods. Based on the comparison (Figure 4 a,b), S-PLM surpassed the performance of the other two PLMs (ARI of ESM2: 0.63 and PromptProtein: 0.46, respectively).

**Figure 4.**
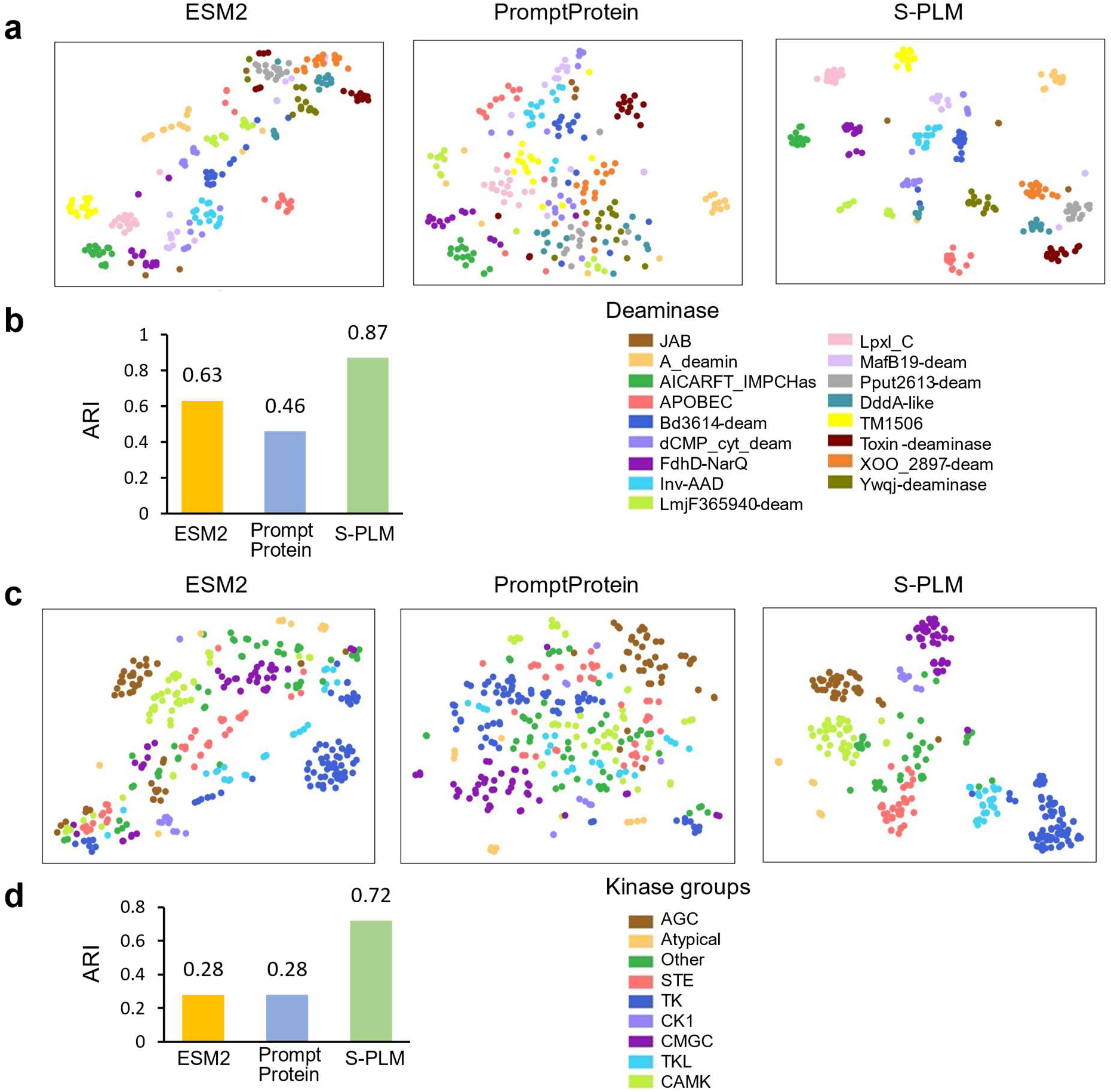
Clustering enzymes via S-PLM. **a**, The t-SNE visualization of protein embeddings from 242 deaminase proteins from sequence representation methods: ESM2, PromptProtein, and S-PLM. **b**, Quantitative benchmarks of the ability of S-PLM to cluster the deaminase proteins for sequence representation methods. **c**, The t-SNE visualization of protein embeddings from human kinase groups via pretrained PLM models: ESM2, PromptProtein, and S-PLM. Different kinase groups are distinguished by different colors. **d**, Quantitative benchmarks of the ability of S-PLM to cluster the human kinases compared with other methods mentioned in c. For both types of enzymes, ARI was computed by comparing K-Means clustering assignments with enzyme labels.

We also studied another enzyme group, kinases, which facilitate the transfer of phosphate groups to proteins in a critical process known as phosphorylation^34^. We extracted 336 kinases, categorized into 9 kinase groups, along with their respective kinase domain sequences from GPS5.0^35^. Subsequently, sequence embeddings were generated for each kinase using their corresponding kinase domain sequences. For comparison, we also obtained sequence embeddings from two other PLMs, ESM2 (ESM2-t33_650M_UR50D) and PromptProtein. From the t-SNE plots (Figure 4c) and the ARI comparison (Figure 4d), the sequence embeddings produced by our S-PLM showed superior division of kinase groups, with a significantly higher ARI (0.72) compared to ESM2 (0.28) and PromptProtein (0.28). This is likely because the S-PLM model incorporates protein structure information essential for characterizing kinase groups. Taken together, S-PLM provides effective sequence representation for enzyme clustering.

### S-PLM outperforms ESM2 for protein fold and enzyme reaction classification

We evaluated S-PLM for supervised protein fold and enzyme reaction classification, comparing the results with the base ESM2 encoder. The same classification layer was applied on top of each encoder, respectively. The comparison comprises two scenarios. In the first scenario, we treated both S-PLM and ESM2 as protein representation models, freezing all parameters of each encoder and allowing only the classification layers to be trainable. This approach ensures a fair comparison, as the trainable layers in both S-PLM and ESM2 have the same architecture. In the second scenario, we fine-tuned the top-*K* layers of both the S-PLM and ESM2 models, allowing the encoders to be trainable along with the classification layers. It is important to note that, given the different architectures of S-PLM and ESM2, achieving absolute fairness in this comparison is impossible, as the optimal configurations for both models may differ. Consequently, to align more closely with ESM2, all configurations were preselected based on ESM2’s optimal performance, including the experiments in the first scenario. For scenario two, the key hyperparameter of ESM2 is *K*, (i.e., how many top Transformer layers are trainable). For fold classification, we tried *K* = 5,6,7, and for enzyme reaction, we tried *K* = 1,2. Details regarding the model configurations can be found in Table S1 (Supporting Information).

Table 1 shows that the proposed S-PLM model consistently outperformed the base ESM2 models for all the classification tasks when all encoder layers were frozen (ESM2-fix *vs*. S-PLM-fix). When an equivalent number of Transformer layers were fine-tuned (ESM2-finetune top-*K* vs. S-PLM-finetune top-*K*), S-PLM-finetune outperformed in all tasks except for family classification. Despite a slight 0.2% decrease in family classification, S-PLM-finetune demonstrated a notable 2–3% improvement in superfamily and fold predictions. This finding highlights the significance of structural information in fold and superfamily predictions, whereas sequence information prevails in family prediction. We noted that the superior performance of S-PLM-finetune does not stem from more trainable parameters compared to ESM2-finetune. For instance, S-PLM-finetune top5 (116M) has fewer parameters than ESM2-finetune top6 (120M), yet it outperformed in fold (37.74% vs. 36.63%) and superfamily (77.59% vs. 75.84%) predictions. In enzyme reaction classification, our model achieved 86.71% test accuracy, outperforming ESM2-finetune top2 (84.50%) with fewer parameters (23M vs. 40M). Taken together, these results support the benefits of integrating structural information into our S-PLM model, particularly for improving predictions in protein fold and enzyme reaction classifications, where structure features are crucial.

**Table 1:**
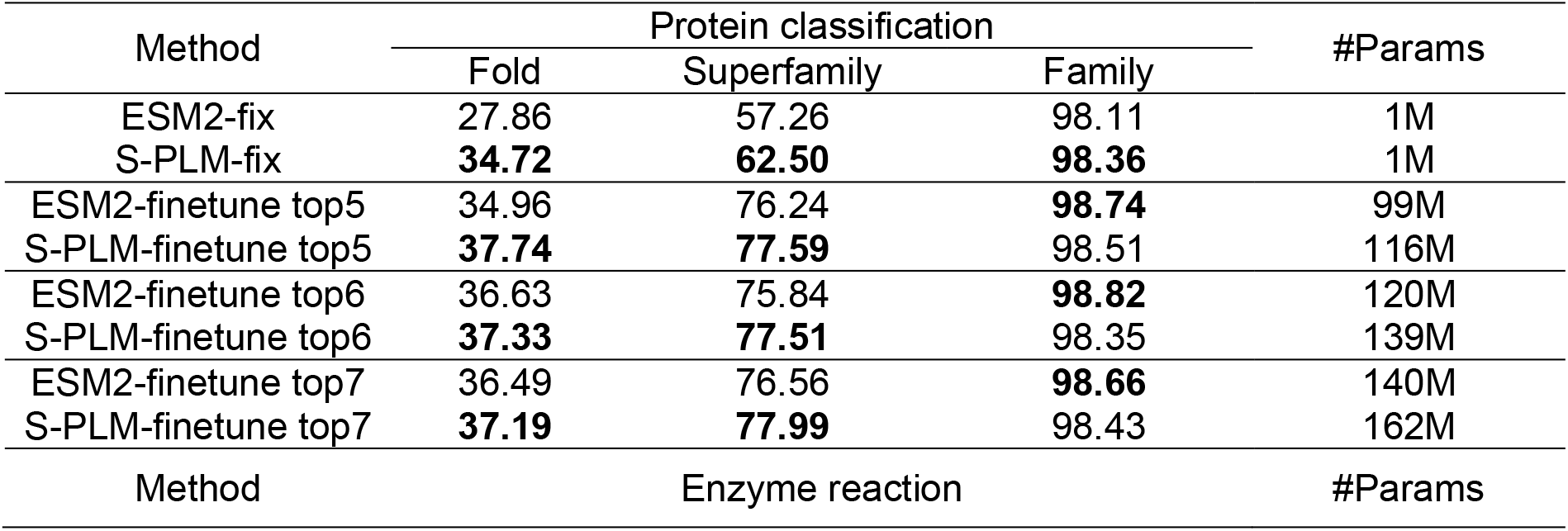
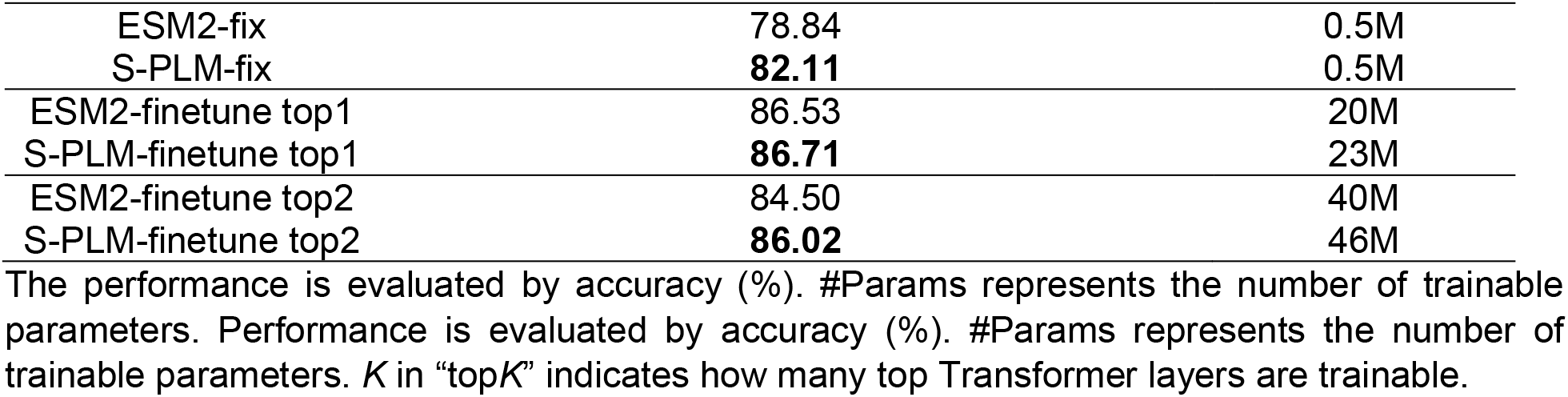
Comparison between S-PLM and ESM2 for protein fold and enzyme reaction classification.

| Method | Protein classification |  |  | #Params |
| --- | --- | --- | --- | --- |
|  | Fold | Superfamily | Family |  |
| ESM2-fix | 27.86 | 57.26 | 98.11 | 1M |
| S-PLM-fix | **34.72** | **62.50** | **98.36** | 1M |
| ESM2-finetune top5 | 34.96 | 76.24 | **98.74** | 99M |
| S-PLM-finetune top5 | **37.74** | **77.59** | 98.51 | 116M |
| ESM2-finetune top6 | 36.63 | 75.84 | **98.82** | 120M |
| S-PLM-finetune top6 | **37.33** | **77.51** | 98.35 | 139M |
| ESM2-finetune top7 | 36.49 | 76.56 | **98.66** | 140M |
| S-PLM-finetune top7 | **37.19** | **77.99** | 98.43 | 162M |
| Method | Enzyme reaction |  |  | #Params |

|  |  |  |
| --- | --- | --- |
| ESM2-fix | 78.84 | 0.5M |
| S-PLM-fix | **82.11** | 0.5M |
| ESM2-finetune top1 | 86.53 | 20M |
| S-PLM-finetune top1 | **86.71** | 23M |
| ESM2-finetune top2 | 84.50 | 40M |
| S-PLM-finetune top2 | **86.02** | 46M |

The performance is evaluated by accuracy (%). #Params represents the number of trainable parameters. Performance is evaluated by accuracy (%). #Params represents the number of trainable parameters. $K$ in “top $K$ ” indicates how many top Transformer layers are trainable.

**Table 2:** Comparing different lightweight tuning methods for sequence encoders of S-PLM on GO, EC, and SS prediction with their key model descriptions.

| Method | GO (F<sub>max</sub>) |  |  | #Params | Sequence encoder layers | Classificati<br>on layer |
| --- | --- | --- | --- | --- | --- | --- |
|  | BP | MF | CC |  |  |  |
| S-PLM-finetune | 0.470 | 0.674 | 0.460 | 42M,40M,40M | finetune top 2 layers,<br>freeze adapter | mean<br>pooling<br>over all<br>residues |
| ESM2-finetune | 0.473 | 0.677 | 0.450 |  |  |  |
| S-PLM-adapter | **0.495** | **0.685** | **0.484** | 55M,53M,53M | adapter tuning<br>top 16 layers |  |
| ESM2-adapter | 0.493 | 0.683 | 0.472 |  |  |  |
| S-PLM-LoRA | 0.480 | 0.668 | 0.472 | 3M,1M,0.7M | LoRA on top 16<br>layers, $r = 2, \alpha = 8$ | |
| ESM2-LoRA | 0.478 | 0.663 | 0.474 |  |  |  |
| Method | EC (F<sub>max</sub>) |  |  | #Params | Sequence encoder<br>layers | Classificati<br>on layer |
| S-PLM-finetune | 0.885 |  |  | 40M | finetune top 2 layers,<br>freeze adapter | mean<br>pooling<br>over all<br>residues |
| ESM2-finetune | 0.888 |  |  |  |  |  |
| S-PLM-adapter | 0.875 |  |  | 53M | adapter tuning<br>top 16 layers |  |
| ESM2-adapter | 0.866 |  |  |  |  |  |
| S-PLM-LoRA | **0.888** | | | 1M | LoRA on top 33<br>layers, $r = 2, \alpha = 8$ | |
| ESM2-LoRA | 0.864 |  |  |  |  |  |

| Method | SS (Accuracy%) | #Params | Sequence encoder layers | Classification layer |
| --- | --- | --- | --- | --- |
| S-PLM-finetune | **87.48** | 92M | finetune top 2 layers and all adapter layers | MLP (1280, 100, 3) on each residue |
| ESM2-finetune | 87.21 |  |  |  |
| S-PLM-adapter | 87.40 | 52M | adapter tuning top 16 layers |  |
| ESM2-adapter | 86.73 |  |  |  |
| S-PLM-LoRA | 85.72 | 0.8M | LoRA on top 33 layers, $r = 2, \alpha = 8$ | |
| ESM2-LoRA | 87.14 |  |  |  |

### Lightweight tuning strategies help S-PLM achieve state-of-the-art performance in supervised protein predictions

To adapt S-PLM for various specific supervised protein prediction tasks, we developed lightweight tuning strategies, including fine-tuning top layers, adapter tuning (Figure 1c), and LoRA (Figure 1d), on the sequence encoder of S-PLM. We benchmarked S-PLM with these lightweight tuning strategies on the GO term, EC number, and secondary structure predictions with state-of-the-art (SOTA) methods. We employed the same training, validation, and test sets for all these tasks, and the only differences are our models and the training strategies. The GO term and EC number predictions served as protein-level supervised classification tasks, which are evaluated in terms of the F_max_, while the secondary structure (SS) prediction served as the residue-level supervised classification task evaluated by accuracy (%). The comparison results are shown in Figure 5. We use ‘our model’ to represent the performance of S-PLM with the most effective lightweight tuning strategy tailored to each task. Comprehensive results across all three training strategies, along with detailed model descriptions, are provided in Table 2. For GO and EC number predictions, five methods—GVP^14^, GearNet^36^, GearNet-Edge^36^, CoupleNet^15^, and ESM-GearNet^14^—that use both sequence and structure inputs, as well as sequence-only method ESM2 (650M)^14^, were considered for comparison. For the SS task, sequence-only methods, TAPE (Transformer)^37^, ProteinBERT^12^, ESM-1b^38^, and DML^39^ were considered for comparison. Their results were taken directly from their respective publications.

**Figure 5.**
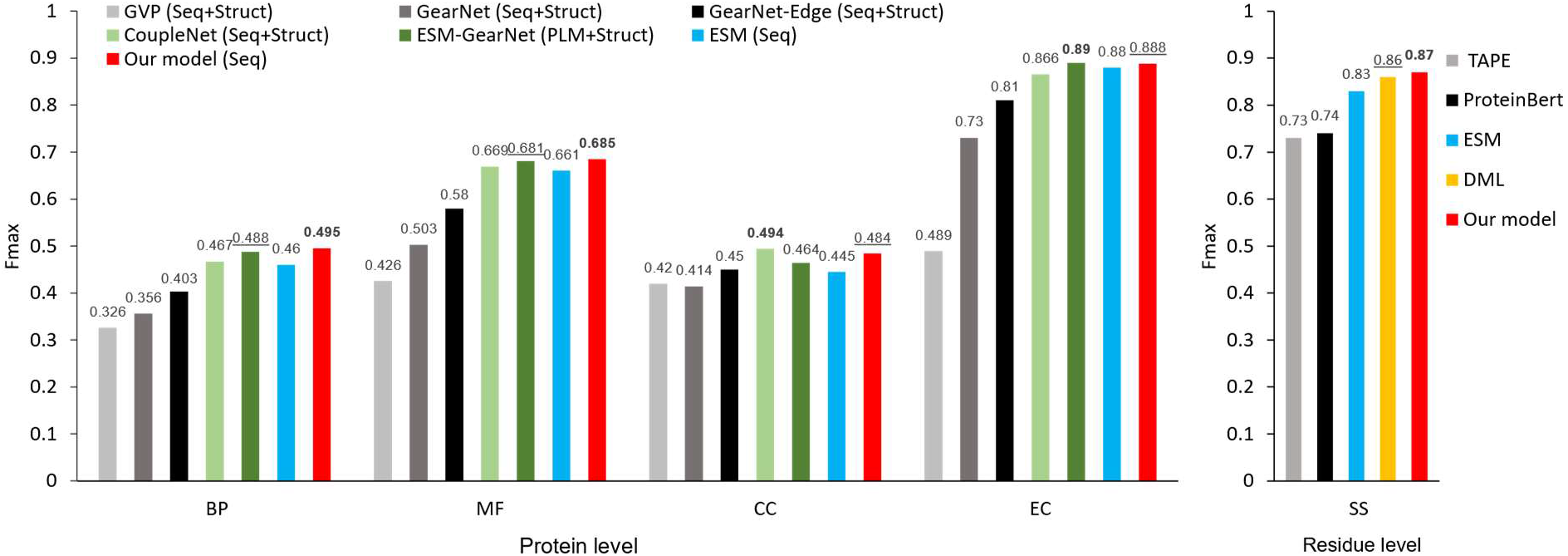
Benchmarking on GO, EC, and SS prediction. Left: Protein level tasks (BP, MF, CC, and EC); Right: Residue level task (SS). “Seq,” “Struct,” and “PLM” indicate the usage of protein sequence, structure, and PLM information in the model, respectively. Five methods that use both sequence and structure inputs (GVP^14^, GearNet^36^, GearNet-Edge^36^, CoupleNet^15^, and ESM-GearNet^14^), as well as sequence-only method ESM2 (650M)^14^ were considered for comparison on GO and EC tasks. For the SS task, sequence-only methods, TAPE (Transformer)^37^, ProteinBERT^12^, ESM-1b^38^, and DML^39^ were considered for comparison. The best and the second-best results for each task are shown in bold and underlined symbols, respectively.

Upon comparison, our model demonstrated SOTA performance in GO-BP (F_max_: 0.495), GO-MF (F_max_: 0.686), and SS (F_max_: 87.48) tasks, while exhibiting suboptimal performance in GO-CC (CoupleNet: 0.494 vs. our model: 0.484 of F_max_) and for EC number prediction (ESM-GearNet: 0.890 vs. our model: 0.888 of F_max_). Among the methods under comparison, GVP and GearNet are designed to capture the invariant and equivariant features of protein structure, whereas GearNet-Edge is a variant of GearNet enhanced with an edge message passing layer. CoupleNet requires the integration of sequence and structure information by deeply coupling them, and ESM-GearNet is a recently proposed variant of GearNet that combines the representation of sequence from PLM with the representation of structure and integrates the two modalities through various fusion schemes, where the results with the best fusion scheme were reported in the comparison. In contrast to these methods, our model relies solely on protein sequences. Remarkably, it achieves compatible results (compared to CoupleNet and ESM-GearNet) or even superior results (compared to GVP, GearNet, and GearNet-Edge), showcasing the broader practical utility of our model.

Additionally, by employing the same lightweight tuning strategies on the base ESM2 encoder with an equal number of trainable parameters, we established a fair comparison between our model and its corresponding base model. The results are presented in Table 2. In most (10/15) of the one-to-one comparisons, our S-PLM encoder outperformed its base ESM2 encoder. Although we could not surpass ESM2 across all tasks under all training strategies, our S-PLM model consistently achieved the best performance overall for each individual task. Overall, our additional Struct-Adapter on ESM2, along with the new pretraining process, did not diminish ESM2’s original sequence representation capability but rather led to superior performance across all tasks.

## Discussion and conclusion

In this work, we proposed S-PLM, a structure-aware protein language model pretrained via contrastive learning between sequence and 3D structure of proteins. Distinguishing from joint-embedding-based methods, S-PLM independently encodes protein sequences and 3D structures. This unique feature allows S-PLM to make predictions using only protein sequences, reducing dependence on the protein’s 3D structure. This aspect is particularly important when obtaining reliable protein structure is difficult or time-consuming. By employing a Struct-Adapter module on the pretrained ESM2 model, our method generated sequence representations that seamlessly incorporate 3D structure information while maintaining the original sequence representation capabilities of ESM2. Importantly, this adapter is extensible; in the future, if new protein attributes need to be included, such as protein function and protein-protein interactions, the existing ESM2 and structure adapter can remain unchanged. Simply adding a new adapter and training on the updated data will suffice (e.g., Adapter 1 and Adapter 2 in Figure 1c).

The results of our study highlight the efficacy of S-PLM across a range of downstream use cases. First, S-PLM demonstrates the ability to align sequence and structure embeddings of the same protein effectively while keeping embeddings from other proteins further apart. Secondly, compared to other PLMs that only use sequences, S-PLM shows impressive awareness of protein structures, as evidenced by its superior performance in clustering CATH domains and enzyme clustering tasks. Finally, together with S-PLM model, we developed a library of lightweight training strategies that can be applied to train S-PLM. Even without inputting protein structure, S-PLM achieves competitive performance in protein supervised prediction tasks, comparable to the SOTA methods that require both sequence and structure inputs. These findings collectively highlight the potential of S-PLM, together with its lightweight tuning toolbox, as an alternative PLM for protein analysis and prediction tasks.

We also observed that different tuning strategies exhibited varying performance across tasks (Table 2). This can be attributed to several factors. First, the number of trainable parameters plays a role, with fine-tuning top layers generally providing more trainable parameters than adapter tuning, and LoRA having the fewest trainable parameters. However, the extent of adaptation to which the base model can be modified is also crucial. Despite having fewer trainable parameters, LoRA’s ability to influence more layers of the base model may enable it to capture complex patterns better suited for certain tasks that diverge significantly from the pretraining objective. Additionally, the nature of the tasks themselves likely contributes to the performance differences. Some tasks may benefit more from fine-grained adaptation of the lower layers, while others might require more substantial changes to the higher-level representations. The diverse lightweight training strategies we provide offer flexibility in tailoring the number of trainable parameters and the extent of adaptation, potentially enabling SOTA performance on a wide range of downstream tasks. Moreover, our S-PLM model’s additional Struct-Adapter module, which infuses structural information into the pretrained sequence representations, introduces an inherent capacity advantage over the base ESM2 model when tuning the same number of parameters. This structural awareness may prove advantageous for tasks that rely heavily on understanding the intricate relationships between sequence and structure.

Although we have achieved encouraging results by training on the AlphaFold2-SwissProt database (0.5M), potential improvement is possible by expanding our training to encompass more comprehensive protein structure repositories. Our current adapter-tunning approach lays a solid foundation for future continuous lifelong learning. By continuously refining and expanding the training data, we aim to imbue S-PLM with a broad range of protein structures across diverse biological contexts. This iterative process of data augmentation and model refinement holds promise for our method to push the boundaries of sequence-based protein representation learning.

## Methods

### 1 Sequence encoder

Our sequence encoder was developed based on the pretrained ESM2 model^5^. Given the constraints of computational resources and model capacity, we selected ESM2-t33_650M_UR50D as our base PLM model, which has 650 million parameters. In particular, we first tokenized the input protein sequence by one-hot encoding for each amino acid and then applied 33 layers of Transformer encoders. The embedding dimension for each position was 1280. In this procedure, a BEGIN token (“<cls>“) and END token (“<eos>“) were added to the sequence and went through the Transformer together with the amino acid tokens, and a “<pad>” token was used for padding sequences. Through the Transformer layers, the output was a tensor of 1280D vector per residue, with the embedding of the BEGIN and END tokens as well as the embeddings for padding sequences. The embeddings for each residue were used for residue-level representations of a protein for downstream tasks. The average of per-residue embeddings, excluding the padding tokens, was used for protein-level embedding for the contrastive-learning training and downstream tasks. Then, two projector layers were applied to the protein-level embedding that transformed the dimension into the final output protein-level embedding, which was 256D. The projector layers were two linear layers.

### 2 Structure encoder

We utilized the contact map to represent the 3D protein structure because it contains complete protein structure information, possesses inherent invariance capabilities, and is straightforward for implementation. Therefore, our structure encoder is specifically designed to encode the protein contact maps. Because the contact map representation resembles an image, any widely adopted networks for image-related takes can be employed as the encoder. We considered ResNet50^40^, Segment Anything Model (SAM)^41^, and Swin-Transformer^28^ models due to their popularity as image networks and superior performance in various competitions. By evaluating them with pretrained weights on the CATH domains with known protein structures (Figure S2, Supporting Information), we finally applied the Swin-Transformer (swinv2-tiny-patch4-window8-256) because it enables more effective feature extraction from the contact map representation. To meet the requirement of the image network, which expects three input channels, we transformed our contact map into a representation with three channels. We generated the raw contact map by calculating the coordinate distance between the C_α_ atoms for each amino acid for one sequence. In general, a contact map is a binary matrix with a value of 1 if the pairwise distance is within a chosen threshold, indicating contact between the residues; otherwise, its assigned value is 0.

In our case, we applied a distance threshold for each channel and converted the raw contact map into a continuous similarity matrix. Specifically, the distance threshold for each channel was *d*: 22Å, the same value used in AlphaFold2. The value of the similarity matrix ranges from 0 to 1, with 1 indicating the shortest pairwise distance, and 0 indicating the longest distance. The final contact map representation can be formulated as the following:

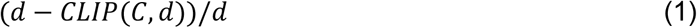

where *C* represents the element of the raw contact map, and CLIP is a function that can change the *C* values higher than *d* into *d*. By averaging the embeddings of the representation layer of Swin-Transformer and excluding the padding representation, we got the protein-level embedding of structure for the contrastive-learning training. Then, we applied two projector layers to the protein-level embedding to transform the dimension into the final output protein-level embedding, which was 256D, the same as the final output protein-level embedding from sequence.

### 3 Multi-view contrastive learning

The objective of contrastive learning in this study is to bring closer the sequence embeddings and structure embeddings from the same protein and further repel all the embeddings from different proteins in the latent space. To achieve this, we applied a multi-view contrastive loss function to the protein-level embeddings obtained from the last projection layer of the sequence and structure encoders. The multi-view contrastive loss function was modified based on the *NT-Xent* (the normalized temperature-scaled cross-entropy loss) in SimCLR^29^. In contrast to the SimCLR paper, in our approach, the positive pair was only defined for sequence embedding 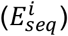 and structure embedding 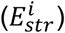 from the same protein *i*. Then the multi-view contrastive loss function for the positive pair 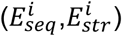 is defined in the following equation :

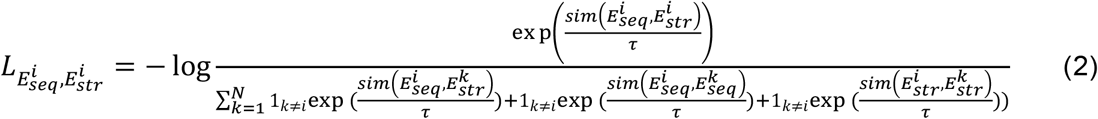

where 1_*k*-≠*i*_ ∈ {0,1} is an indicator function equaling 1 if *k* ≠ *i* and *τ* denotes a temperature parameter; *N* is the batch size. The *sim*(*x, y*) is a function that quantifies the similarity between embeddings *x* and *y*, defined as follows:

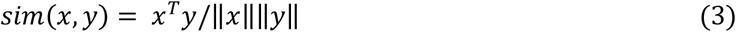

This loss function helps to maximize the alignment of the embeddings for protein *i* in the two views 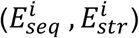 and minimize the alignment between protein *i* and the other proteins (*k* ≠ *i*). The final contrastive loss was calculated for all positive pairs within a mini-batch of samples.

### 4 Preparation of training dataset for contrastive learning and training process

We prepared the training database by obtaining the protein’s amino acid sequences from the Swiss-Prot library and saving them in the FASTA format. The 3D structures of the proteins were obtained from the AlphaFold2 repository. The C_α_-C_α_ contact maps for individual proteins were determined using in-house Python code based on the AlphaFold2 predicted 3D structures. We randomly selected 500,000 proteins from the Swiss-Prot library for training and 41,500 proteins for validation. We did not remove sequences in the validation set that were similar to the ones in the training set. Given the huge size of the Swiss-Prot library (542,378 proteins in total), such similarity is negligible. The training was performed on a single A100 GPU using the computational facility of the University of Missouri. The batch size was 20 due to the GPU RAM limit. S-PLM was trained for more than 20 epochs.

### 5 Lightweight tuning on sequence encoder

The lightweight tuning strategy in this paper refers to training approaches that make specific and often parameter-efficient modifications to a preexisting model, reducing the computational resources and memory needed compared to training a model from fully fine-tuning. We implemented the fine-tuned top layers, LoRA, and adapter tuning on the sequence encoder of S-PLM, as shown in Figure 1, for downstream protein sequence prediction tasks. The details for each strategy are as follows:

Fine-tune top layers: The ESM2 backbone model had 33 total Transformer layers. Here, “fine-tune top layers” refers to fine-tuning only the top *K* ≤ 33 transformer layers and freezing the remainder. Here, *K* is a hyperparameter in our configuration.

LoRA: LoRA refers to low-rank adaptation^23^, which freezes the pretrained model weights and injects trainable rank decomposition matrices into each layer of the Transformer architecture. It can greatly reduce the number of trainable parameters for downstream tasks. For a pretrained weight matrix *W*_0_ ∈ ℝ^*dxk*^, its update Δ*W* can be represented as a low-rank decomposition Δ*W* = *BA*, where *B* ∈ ℝ^*dxr*^, *A* ∈ ℝ^*rxk*^, and the rank *r* ≪ min (*d, k*). During training, *W*_0_ is frozen, *A* and *B* contain trainable parameters. For an input *x*, the LoRA forward pass yields the following:

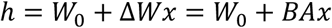

In the implementation of LoRA, Δ*Wx* is scaled by 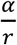, where α is a constant in *r* . In our implementation, the low-rank decomposition matrices were applied to the query, key, value, and output projection matrices in the self-attention module of the top-*K* Transformer layers of ESM2 (Figure 1d). Here, *K*, α, and *r* are hyperparameters in our configuration.

Adapter tuning: Adapter tuning involves integrating adapter modules into the Transformer layer of the ESM2 model. The adapter modules were implemented according to Houlsby’s study^21^. It was positioned twice in one Transformer layer of ESM2: after the self-attention projection and after the two feed-forward. Each adapter module consisted of a bottleneck structure and a skip connection. The bottleneck structure compressed the input data into a reduced-dimensional space and then reconstructed the data to restore it to the original input dimension. The bottleneck structure enabled the adapter module to have few parameters relative to the attention and feed-forward layers in the original Transformer. The integration of the ESM2 model with the structure-aware adapter modules (Struct-Adapter) is illustrated in Figure 1b. Unlike the original adapter tuning^21^, which applied the adapter modules into all Transformer layers, we specifically inserted them into the top-*K* Transformer layers of ESM2. In our configuration, *K* is a hyperparameter. Adapter tunning for supervised downstream tasks was implemented by integrating a list of additional paralleled adapters on the Struct-Adapter modules, as shown in Figure 1c. The hyperparameter used and the number of trainable parameters for each strategy for all the downstream tasks are shown in Table 2.

### 6 Data preprocessing for CATH superfamily clustering

We downloaded the sequence data with CATH annotations from the CATH database (release v4_3_0)^31^ and the cath-dataset-nonredundant-S40-v4_3_0 dataset, whose proteins of maximally 40% sequence identity were used. We only kept sequences that had records of known protein structures. We only selected one represented sequence with the longest sequence length from one CATH superfamily. We considered all the three most represented categories of class (1,553 sequences), the five most represented categories of architecture (1,049 sequences), and the five most represented categories of topology (306 sequences) from the CATH hierarchy. The selected CATH sequences are provided in Data 1 (Supporting Information).

### 7 Downstream supervised protein prediction tasks

The downstream supervised protein prediction tasks included four protein-level prediction tasks— protein fold classification, enzyme reaction classification, GO term prediction, and EC number prediction—and one residue-level prediction task, protein secondary structure prediction.

#### 7.1 Fold classification

Protein fold prediction is utilized to accurately determine the fold class label of a given protein. We used the same dataset splits from the study^42^. The dataset comprised 16,712 proteins, classified into 1,195 distinct fold classes. The dataset has three test sets: “Fold,” “Superfamily,” and “Family.” In the “Fold” set, training is conducted, excluding proteins from the same superfamily. In the “Superfamily” set, proteins from the same family are excluded from training. In the “Family” set, proteins belonging to the same family are included in the training process.

#### 7.2 Enzyme reaction classification

The goal of enzyme reaction classification is to determine the class of enzyme-catalyzed reactions for a protein, utilizing all four levels of EC numbers^43^. Following the methodology described in the study^44^, the dataset was split into training, validation, and test sets, comprising 37,428 proteins categorized across 384 four-tiered EC numbers. All proteins had less than 50% sequence similarity across the data splits.

#### 7.3 GO term prediction

The objective of GO term prediction is to ascertain the association of a protein with a specific GO term, including three tasks: biological process (BP), molecular function (MF), and cellular component (CC). Each task is formulated as a multi-label classification problem. We used the same dataset splits from the study^45^, where the test set has up to 95% (30%, 40%, 50%, 70%, and 95%) sequence similarity with the training set.

#### 7.4 EC number prediction

The objective of this task is to identify the 538 distinct third- and fourth-level EC numbers, which characterize the catalytic functions of various proteins in biochemical reactions. It is formulated as a multi-label classification problem. We used the same dataset splits from the paper^45^, where the test set has up to 95% (30%, 40%, 50%, 70%, and 95%) sequence similarity with the training set.

#### 7.5 Secondary structure prediction

This task aims to predict the local structures of protein residues into three secondary structure labels (i.e., coil, strand, or helix). We used the same dataset splits in the reference^38^, adopting Klausen’s dataset^46^ as the training set and the CB513 dataset^47^ as the test set. No two proteins have greater than 25% sequence identity, and the test set has less than 25% sequence identity against the training set. Same as in the reference^38^, we truncated those sequences with more than 1,022 residues by keeping the first 1,022 residues. The statistics of the dataset for each task is shown in Table S2 (Supporting Information). The data can be downloaded from https://github.com/duolinwang/S-PLM/tree/main/SPLM_Data/.

### 8 Implementation of the lightweight tunning tools

The source code of the lightweight tunning tools is available at https://github.com/duolinwang/S-PLM/. Developed primarily using PyTorch (version 12.1), these tools facilitate training and evaluation for the protein prediction tasks described in Methods 7. Users can reproduce the results presented in this paper by retraining with the provided configuration files in the <u>“configs”</u> folder. For customized training with user-specific data, users should incorporate specific data-processing functions into the data.py file to adapt the data format to our model. Protein-level prediction tasks, such as fold and GO tasks, can be referenced for protein-level prediction, while the secondary structure task serves as a reference for residue-level prediction. Additionally, users have the flexibility to select the desired lightweight tuning strategy and modify specific parameters by adjusting the parameters in the configuration files.

## Acknowledgments

D.W. and D.X. are partially supported by the National Institutes of Health (grant R35GM126985). U.A., Q.S., and J.C. would like to thank AI in Medicine (AIM) at the University of Kentucky (NCATS UL1TR001998, NCI P30 CA177558). U.A. and Q.S. would also like to thank the Start-up Fund of the University of Kentucky and Alzheimer’s Association (AARG-23-1144638) for their financial support. Q.S. also acknowledges the financial support of the National Science Foundation (2154996). The computation for this work was partially performed on the high-performance computing infrastructure provided by Research Computing Support Services at the University of Missouri. We also thank the University of Kentucky Center for Computational Sciences and Information Technology Services Research Computing for their support and use of the Morgan Compute Cluster and associated research computing resources. This work also used Delta-GPU at NCSA through allocation CIS230053 from the Advanced Cyberinfrastructure Coordination Ecosystem: Services & Support (ACCESS) program, which is supported by National Science Foundation grants #2138259, #2138286, #2138307, #2137603, and #2138296.

