## supporting information for "S-PLM: Structure-aware Protein Language Model via Contrastive Learning between Sequence and Structure"

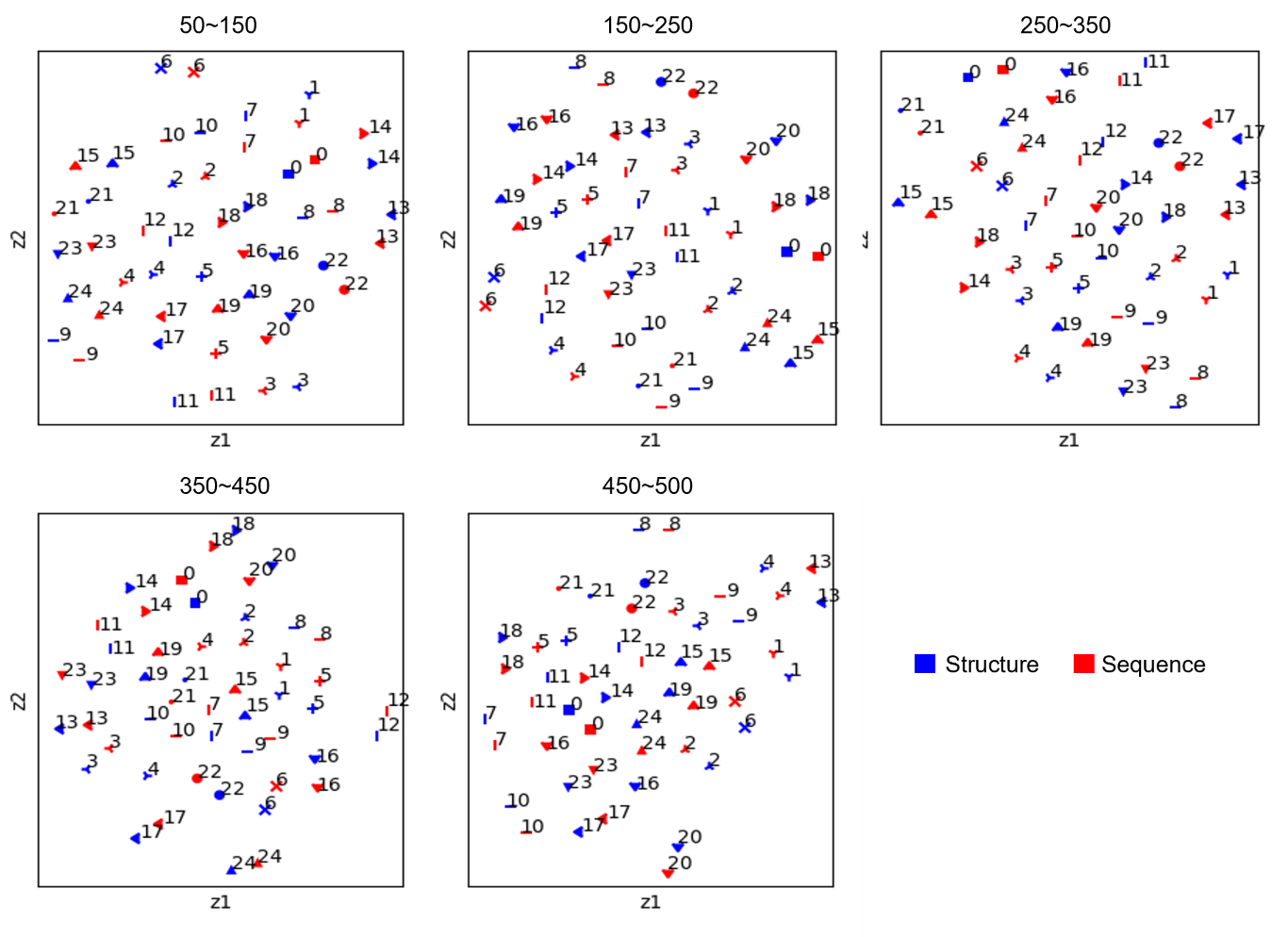

Figure S1. The alignment between structure and sequence embeddings for specific protein sequence length ranges. All the embeddings were generated after model training using contrastive learning. We randomly selected 25 proteins in a sequence length range from the independent test set. A node in blue indicates a structure-based t-SNE embedding for a protein. A node in red indicates a sequence-based t-SNE embedding for a protein. The number beside each node indicates the index of the protein. Different indices and shapes indicate different proteins. The structure-based and sequence-based embeddings from the same protein have the same shape and protein index.

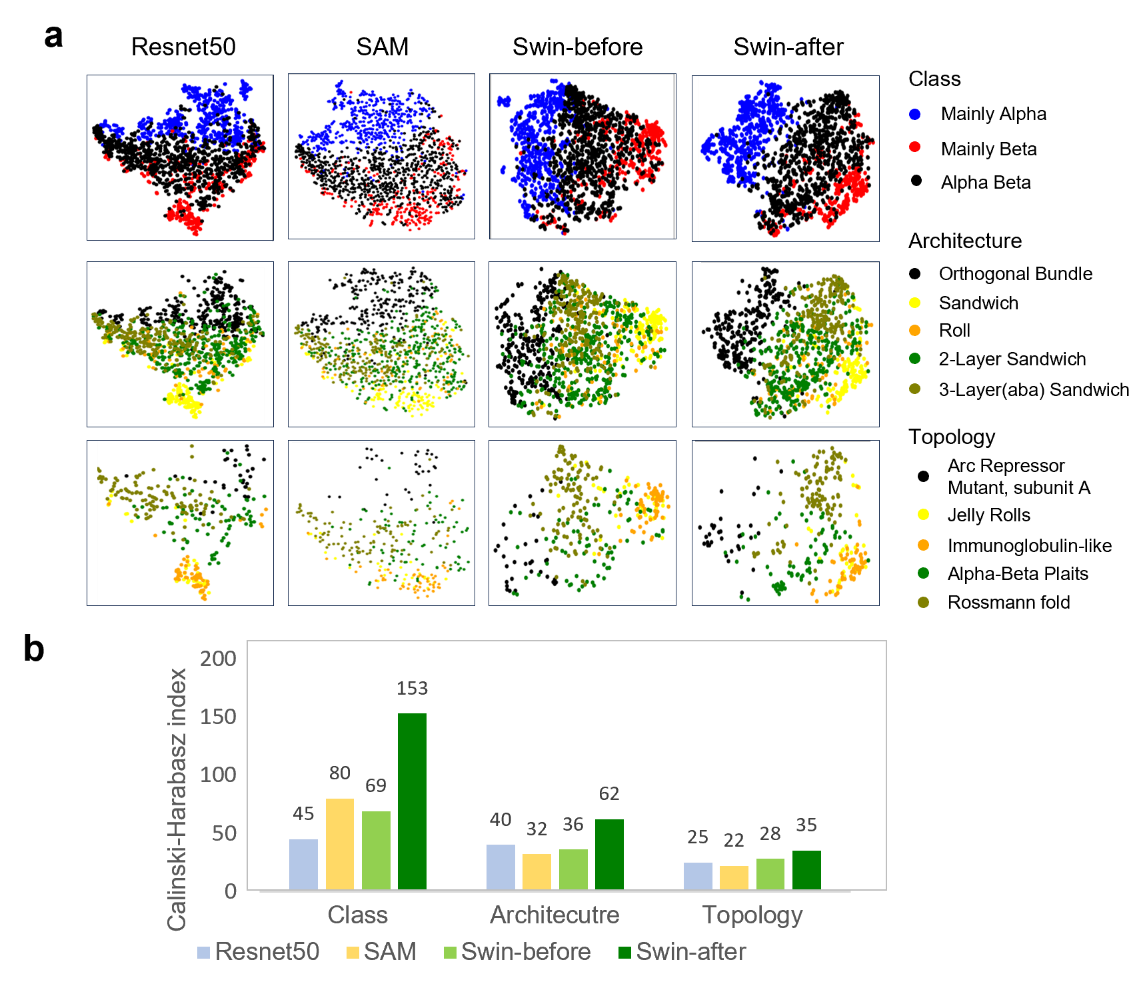

Figure S2. Visualization and benchmark of protein contact map embeddings for three CATH structural hierarchies (Class, Architecture and Topology). **a**, t-SNE visualization of protein contact map embeddings from top-five most represented categories from one hierarchy on contact maps calculated from known PDB proteins. Embeddings under comparison were produced by ResNet50, SAM, Swin-Transformer before (Swin-before) the contrastive learning and Swin-Transformer after (Swin-after) contrastive learning. ResNet50 was pretrained on the imagenet1k_v1 dataset, SAM used pretrained ViT-B encoder, and the Swin-Transformer used the pretrained swinv2-tiny-patch4-window8-256 encoder. **b**, Quantitatively assess the capability of contact map embeddings derived from different structure encoders in clustering CATH structural categories using the Calinski-Harabasz index.

Table S1. Details of S-PLM and ESM2 model configurations for protein fold and enzyme reaction classification.

| **Method** | **Sequence encoder layers** | **Classification layer** | **Optimizer** | **Learning**  **rate** | **# of Epochs** | | |
| --- | --- | --- | --- | --- | --- | --- | --- |
| **Fold classification** | | | | | | | |
| ESM2-fix | freeze ESM2 650M | mean pooping over all residues | Adam | 1e-5 | | 200 | |
| S-PLM-fix | freeze S-PLM |  |  |  |  |  |  |
| ESM2-finetune top 5,6,7 | fine-tune top 5,6,7 layers of Transformer modules |  | Adam | 1e-5 | | 200 | |
| S-PLM-finetune top 5,6,7 | fine-tune top 5,6,7 layers of Transformer-adapter modules |  |  |  |  |  |  |
| **Enzyme reaction** | | | | | | | |
| ESM2-fix | freeze ESM2 650M | mean pooping over all residues | Adam | 1e-3 | | | 80 |
| S-PLM-fix | freeze S-PLM |  |  |  |  |  |  |
| ESM2-finetune top 1 and 2 | fine-tune top 1,2 layers of Transformer modules |  | Adam | 1e-3 | | | 200 |
| S-PLM-finetune top 1 and 2 | fine-tune top 1,2 layers of Transformer-adapter modules |  |  |  |  |  |  |

Table S2: Dataset statistics of downstream supervised protein prediction tasks.

| Dataset | | # of Train | # of Validation | # of Test |
| --- | --- | --- | --- | --- |
| Fold classification | Fold | 12, 312 | 736 | 718 |
|  | Superfamily | 12, 312 | 736 | 1,254 |
|  | Family | 12, 312 | 736 | 1,272 |
| Enzyme reaction | | 29,215 | 2,562 | 5,651 |
| GO term | | 29,898 | 3,322 | 3,415 |
| EC number | | 15,550 | 1,729 | 1,919 |
| Secondary structure | | 8678 | 2170 | 513 |

“#” represents the number of proteins.
